# MicroRNA-mediated control of developmental lymphangiogenesis

**DOI:** 10.1101/555664

**Authors:** Hyun Min Jung, Ciara Hu, Alexandra M. Fister, Andrew E. Davis, Daniel Castranova, Van N. Pham, Lisa M. Price, Brant M. Weinstein

**Author notes:** Correspondence: Brant M. Weinstein, Division of Developmental Biology, Eunice Kennedy Shriver National Institute of Child Health and Human Development, National Institutes of Health, Bethesda, MD, 20892.

## Abstract

The post-transcriptional mechanisms contributing to molecular regulation of developmental lymphangiogenesis and lymphatic network assembly are not well understood. Here, we use high throughput small RNA sequencing to identify miR-204, a highly conserved miRNA dramatically enriched in lymphatic vs. blood endothelial cells, and we demonstrate that this miRNA plays a critical role during lymphatic development. Suppressing miR-204 leads to loss of lymphatic vessel formation, while overproducing miR-204 in lymphatic vessels accelerates lymphatic vessel formation, suggesting a positive role during developmental lymphangiogenesis. We also identify the NFATC1 transcription factor as a key conserved target for post-transcriptional regulation by miR-204 during lymphangiogenesis. While miR-204 suppression leads to loss of lymphatics, knocking down its target NFATC1 leads to lymphatic hyperplasia, and the loss of lymphatics in miR-204-deficient animals can be rescued by NFATC1 knockdown. Together, our results highlight a miR-204/NFATC1 molecular regulatory axis required for proper lymphatic development.

## INTRODUCTION

The lymphatic system is important for fluid and protein homeostasis, lipid transport, and immunity. Lymphatic malfunction is linked to many pathologies including lymphedema, cancer metastasis, and cardiovascular disease (Petrova and Koh, 2018, Venero Galanternik et al., 2016, Alitalo, 2011, Stacker et al., 2014, Lim et al., 2013). Primitive lymphatic vessels are derived from lymphatic endothelial cell (LEC) progenitors that transdifferentiate from venous blood endothelial cells (BECs) and subsequently migrate away from the veins to form lymphatic vessels (Sabin, 1902, Hong et al., 2002, Koltowska et al., 2013, Nicenboim et al., 2015). A variety of factors have been identified that are required for proper regulation of lymphangiogenesis during development and disease, including PROX1, SOX18, COUPTFII, VEGFC/VEGFR3, NRP2, CCBE1, CLEC2, YAP/TAZ, and NFATC (Wigle and Oliver, 1999, Dumont et al., 1998, Karkkainen et al., 2004, Francois et al., 2008, Srinivasan et al., 2010, Yuan et al., 2002, Uhrin et al., 2010, Hogan et al., 2009, Kulkarni et al., 2009, Sweet et al., 2015, Bui et al., 2016, Cho et al., 2019). Some of our current knowledge regarding development of the lymphatic system has come from research using the zebrafish, a superb model for studying vertebrate organogenesis with optically clear, externally developing embryos ideal for high-resolution live imaging. The development of the zebrafish lymphatic vascular system is highly conserved and stereotyped (Yaniv et al., 2006, Okuda et al., 2012, Jung et al., 2017, Kuchler et al., 2006), and the availability of transgenic zebrafish expressing fluorescent reporters in lymphatic endothelium makes it straightforward to monitor lymphatic development at the single cell level *in vivo* and visualize even subtle lymphatic defects using optimized high-resolution microscopy technologies (Jung et al., 2016).

MicroRNAs (miRNAs) are important posttranscriptional regulators that play crucial roles in developmental, physiological, and disease-related processes in animals (Gebert and MacRae, 2018). They are approximately 22 nucleotide long noncoding RNAs that guide Argonaute proteins for gene silencing by mRNA degradation and/or translational repression (Jonas and Izaurralde, 2015). Targeting of miRNAs is accomplished by binding of the key nucleotides 2-8 (the “seed” sequence) and additional miRNA sequences to complementary sequences in target mRNAs (Bartel, 2009, Jung et al., 2013, Tay et al., 2008, Lytle et al., 2007). The majority of miRNA target sites are located in the 3’ untranslated regions (UTRs) of mRNAs, although some miRNA targeting also occurs in 5’UTR and coding sequences. MiRNA regulatory networks are thought to confer robustness to biological processes by reinforcing transcription programs and attenuating aberrant transcripts by either switching off or fine-tuning gene expression, helping to buffer against random fluctuations in transcript copy number (Ebert and Sharp, 2012, Staton et al., 2011, Choi et al., 2007). Although we know a great deal about the roles of protein-coding genes during lymphangiogenesis, we still have limited insight into how post-transcriptional mechanisms regulate lymphatic development. Profiling of miRNAs in human lymphatic endothelial cells (LECs) and blood endothelial cells (BECs) has identified BEC-specific miRNAs such as miR-31 and miR-181a that target Prox1 and prevent lymphatic specification in BECs (Pedrioli et al., 2010, Kazenwadel et al., 2010, Dunworth et al., 2014). These studies suggest some miRNAs can help BECs retain their identity by negatively regulating lymphatic development. Manipulation of endothelial miRNAs has also been shown to result in defective lymphatic development (Kontarakis et al., 2018, Nicoli et al., 2012, Chen et al., 2016). Although these studies have begun to shed light on the role of miRNAs during lymphangiogenesis, our understanding of the role lymphatic miRNAs play during lymphatic vessel development is still limited.

Here, we characterize miR-204, a highly conserved lymphatic-enriched miRNA isolated via small RNA sequencing of human endothelial cells, and demonstrate its critical function during lymphjangiogenesis *in vivo* using the zebrafish. MiRNA-204-deficient zebrafish display severe defects in lymphatic vessel formation, while excess mir-204 expression in endothelium drives precocious lymphangiogenesis. We identify NFATC1 as a conserved mir-204 target in both human lymphatic endothelial cells and in the zebrafish, with loss of mir-204-mediated silencing resulting in increased NFATC1 transcript levels. As in mammalian lymphatics, attenuating nfatc1 in the zebrafish promotes abnormal lymphatic expansion, and suppressing nfatc1 rescues lymphatic development in mir-204-deficient zebrafish. Our results thus identify a miR-204/NFATC molecular pathway critical for lymphatic development.

## RESULTS

### miR-204 is enriched in human and zebrafish lymphatic endothelial cells

We performed small RNA sequencing on human lymphatic endothelial cells (LEC) and human blood endothelial cells (BEC) to identify miRNAs enriched in LECs compared to BECs. We used triplicate samples of total RNA isolated from human dermal lymphatic microvascular endothelial cells (HMVEC-dLy, representing LEC) and human umbilical vein endothelial cells (HUVEC, representing BEC) for sequencing (**Figure 1a**). Prior to sequencing, we used TaqMan quantitative RT-PCR to verify that HMVEC-dLy and HUVEC are enriched for markers representing lymphatic or blood vessel identity, respectively. We tested the expression of lymphatic vascular markers PROX1, FLT4 (also known as Vegfr3), and PDPN, and blood vascular markers NR2F2 (also known as COUP-TFII), KDR (also known as VEGFR2), CDH5 (also known as VE-Cadherin), and EGFL7. HMVEC-dLy express higher levels of PROX1, FLT4, and PDPN, while HUVEC showed enrichment for NR2F2, KDR, CDH5, and EGFL7, showing that these cell types appropriately express genes representative of lymphatic or blood vessel identity (**Figure 1-figure supplement 1**). We performed small RNA sequencing collecting a total of ~10 million reads from each sample and aligned using the miRbase v22 (Kozomara and Griffiths-Jones, 2014). We excluded from further analysis any miRNAs that were represented by less than 10 reads in three or more of the six (two triplicate) sequenced samples, resulting in 445 annotated miRNAs. 98 of these miRNAs showed a significant difference between the LEC and BEC samples (p < 0.01, false discovery rate (FDR) < 0.01). The normalized sequencing data have been submitted to the Gene Expression Omnibus repository with accession number GSE126679. Of these, we identified 30 miRNAs highly enriched in LECs (fold change (FC) >4) and 20 miRNAs enriched in BECs (FC >4) (**Figure 1a, Figure 1-source data 1**). Among the 30 LEC-enriched miRNAs, miR-204-5p was by far the most highly enriched in LEC, with 105-fold higher representation in the LEC sequences compared to BEC (**Figure 1b**). Using the TaqMan miRNA qPCR assay, we confirmed that miR-204-5p is very highly enriched in LEC compared to BEC, in contrast to previously reported vascular miRNAs miR-126 and miR-31 that are more highly enriched in BEC (**Figure 1c**).

**Figure 1.**
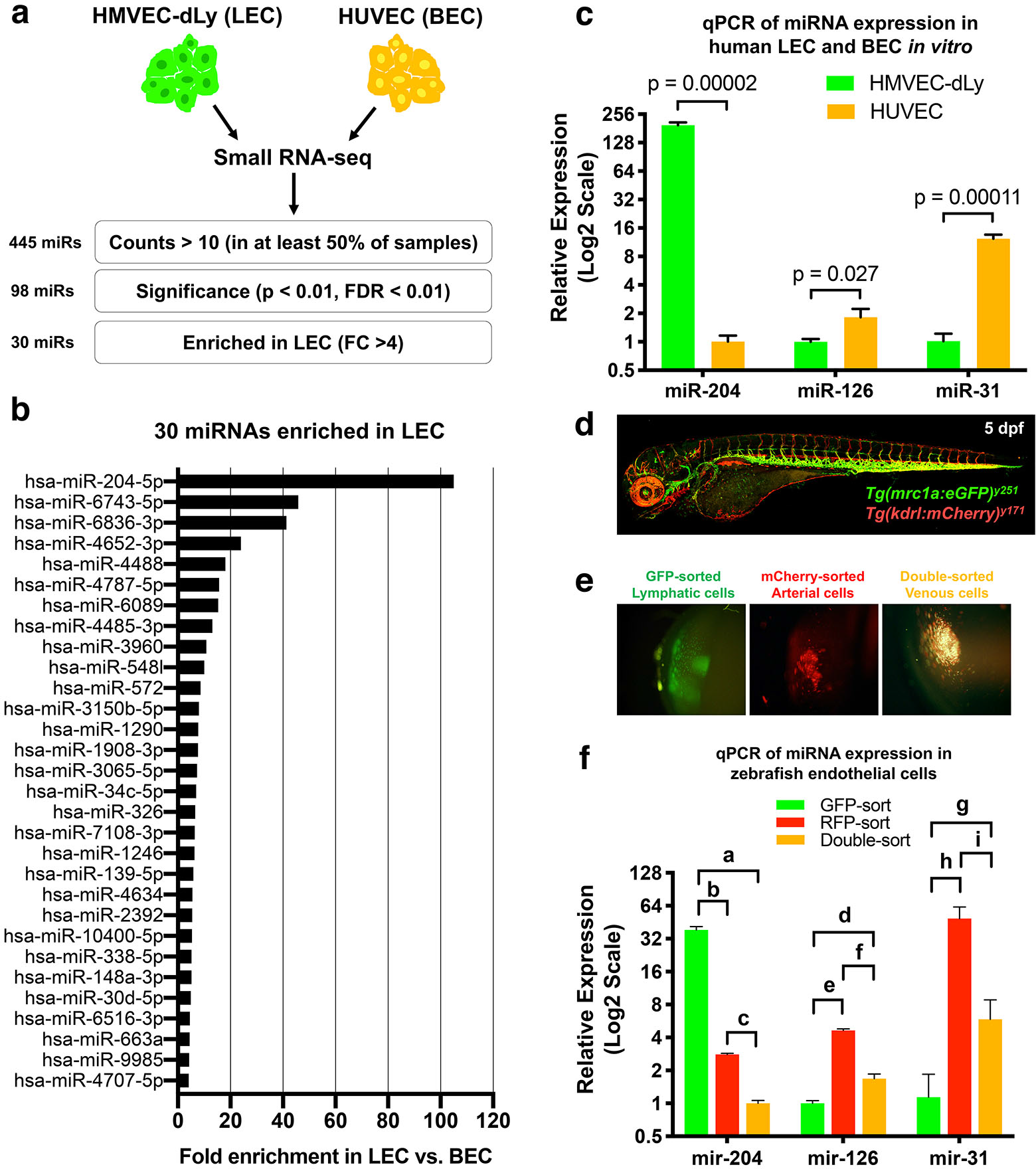
Identification of lymphatic miRNAs enriched in human and zebrafish lymphatic endothelial cells. **(a)**Schematic diagram of the workflow for small RNA sequencing from lymphatic (HMVEC-dLy) and blood (HUVEC) endothelial cells and selection of microRNAs enriched in lymphatic endothelial cells. **(b)**Relative fold enrichment of the 22 most highly enriched miRNAs in LEC versus BEC small RNA sequence data (average of triplicate samples from each group). **(c)**Quantitative TaqMan RT-PCR measurement of the relative expression of three different miRNAs in HMVEC-dLy (LEC) and HUVEC (BEC). Levels of mir-204 are normalized to HUVEC (BEC) levels, while levels of mir-126 and mir-31 are normalized to HMVEC-dLy (LEC) levels. Three biological replicates were analyzed. **(d)**Confocal image of a 5 dpf *Tg(mrc1a:eGFP)*^*y251*^, *Tg(kdrl:mCherry)*^*y171*^ double-transgenic larva (lateral view, rostral to the left). **(e)**Confocal images of lymphatic (GFP-positive), arterial (mCherry-positive), and venous (GFP and mCherry double-positive) endothelial cell pellets isolated from dissociated 5 dpf transgenic animals such as that in panel d by Fluorescence Activated Cell Sorting (FACS). **(f)**Quantitative TaqMan RT-PCR measurement of the relative expression of mature mir-204, mir-126, and mir-31 in FACS-sorted zebrafish endothelial cells. FACS-sorted cells from ~1000 5 dpf larvae were used and three technical replicates were analyzed. Levels of mir-204 are normalized to venous (GFP and mCherry double-positive) levels, while levels of mir-126 and mir-31 are normalized to lymphatic (GFP-positive) levels. P-values (a = 0.000015, b = 0.000018, c = 0.000003, d = 0.0027, e = 0.000004, f = 0.000029, g = 0.055, h = 0.0036, i = 0.0057). All graphs are analyzed by t-test and the mean ± standard deviation (SD) is shown.

We used the zebrafish as an *in vivo* system to examine the role of mir-204 during lymphatic development, beginning by testing whether zebrafish mir-204 is also enriched in LEC. Zebrafish have a conserved, highly stereotyped developing lymphatic vascular network that is readily visualized using transgenic reporter lines (Okuda et al., 2012, Jung et al., 2017) (**Figure 1-figure supplement 2a,b**). We used 5 day post-fertilization (dpf) *Tg(mrc1a:eGFP)*^*y251*^, *Tg(kdrl:mCherry)*^*y171*^ double-transgenic larvae with EGFP-positive lymphatic EC, mCherry-positive arterial EC, and EGFP and mCherry double-positive venous EC to isolate each of these endothelial cell populations by fluorescence activated cell sorting (FACS) (**Figure 1d**). Transgenic larvae were dissociated into single cells and subjected to FACS sorting to obtain EGFP-sorted lymphatic cells, mCherry-sorted arterial cells, and double-sorted venous cells (**Figure 1e**). The EGFP-sorted lymphatic cells showed strong expression of lymphatic markers lyve1b and prox1a, while arterial and venous endothelial cells expressed relatively low levels of these transcripts (**Figure 1-figure supplement 2c**). In contrast, blood vascular markers kdrl and cdh5 were more abundant in mCherry-sorted arterial and double-sorted venous cells compared to EGFP-sorted lymphatic cells (**Figure 1-figure supplement 2d**). Using these sorted endothelial cell populations we were able to show that zebrafish mir-204 is highly enriched in EGFP-sorted lymphatic cells, while the blood endothelial miRNAs mir-126 and mir-31 are more enriched in mCherry-positive or double-positive BECs (**Figure 1f**). These data show that miR-204 is a highly conserved miRNA enriched in the lymphatic endothelium in both humans and zebrafish.

### Developmental lymphangiogenesis is suppressed by miR-204 deficiency

Human miR-204 is located in the sixth intron of TRPM3 (Transient Receptor Potential Cation Channel Subfamily M Member 3), and the mature miRNA is 100% conserved amongst a variety of vertebrate species (**Figure 2-figure supplement 1a**). The zebrafish genome contains three paralogues of mir-204 (http://miRbase.org). As in the human genome, one mir-204 is found in intron 5 of the zebrafish *trpm3* gene (*mir-204-1*) (**Figure 2-figure supplement 1a**). However, additional zebrafish mir-204 sequences are also found in intron 5 of *trpm1a* and in intron 4 of *trpm1b* (*mir-204-2* and *mir-204-3*, respectively; **Figure 2-figure supplement 1b**). The single human *TRPM1* gene contains a closely related paralogue miRNA, miR-211, in intron 6. Although each of the three copies of mir-204 in the zebrafish has unique precursor sequences, their mature mir-204 sequences are 100% identical (**Figure 2a**). To determine the function of mir-204 in lymphatic vessel formation during early development, we began by using morpholino (MO) antisense oligomers to target and block the function of endogenous mir-204 in zebrafish. Knocking down miRNA function using MOs is an excellent targeting strategy for miRNAs because unlike protein-coding genes the functional products of miRNA genes are RNAs (Flynt et al., 2017). Four different MOs were designed to use different strategies to block mir-204 function. The pan-204 MO targets the mature mir-204 sequence and knocks down all mir-204s. MiRNA precursor hairpin structures must be properly cleaved by Dicer to permit maturation of functional miRNAs (Park et al., 2011, Flynt et al., 2017, Kloosterman et al., 2007). We designed specific MOs targeting the dicer-cleavage sites of *mir-204-1*, *mir-204-2*, and *mir-204-3*, respectively, to individually block the maturation of each of the three precursor mir-204 sequences (**Figure 2a**). Injection of 0.5 ng of pan-204 MO led to highly efficient suppression of mature mir-204 levels (**Figure 2b**). At the 0.5 ng dose there were no noticeable morphological anomalies, although slightly higher doses (0.75 ng) resulted in some pericardial edema, jaw defects, mild microcephaly and smaller eye development (**Figure 2c**), so we used the 0.5 ng dose for all experiments with the pan-204 MO. We examined whether mir-204 knockdown affects lymphatic vessel development by observing the parachordal line at 3 dpf (**Figure 2d,e-i**) and the thoracic duct at 5 dpf (**Figure 2d,j-n**). In comparison to control MO injected animals that form a parachordal line (a cord of lymphatic progenitors found at 3 dpf along the trunk midline) pan-204 MO-injected animals displayed loss of the parachordal line (**Figure 2e-i**). In normally developing embryos, some LECs from the parachordal line subsequently migrate ventrally to form the thoracic duct just below the dorsal aorta by 5 dpf (**Figure 2j,k**). Animals injected with the pan-204 MO also displayed loss of the thoracic duct (**Figure 2l-n**).

**Figure 2.**
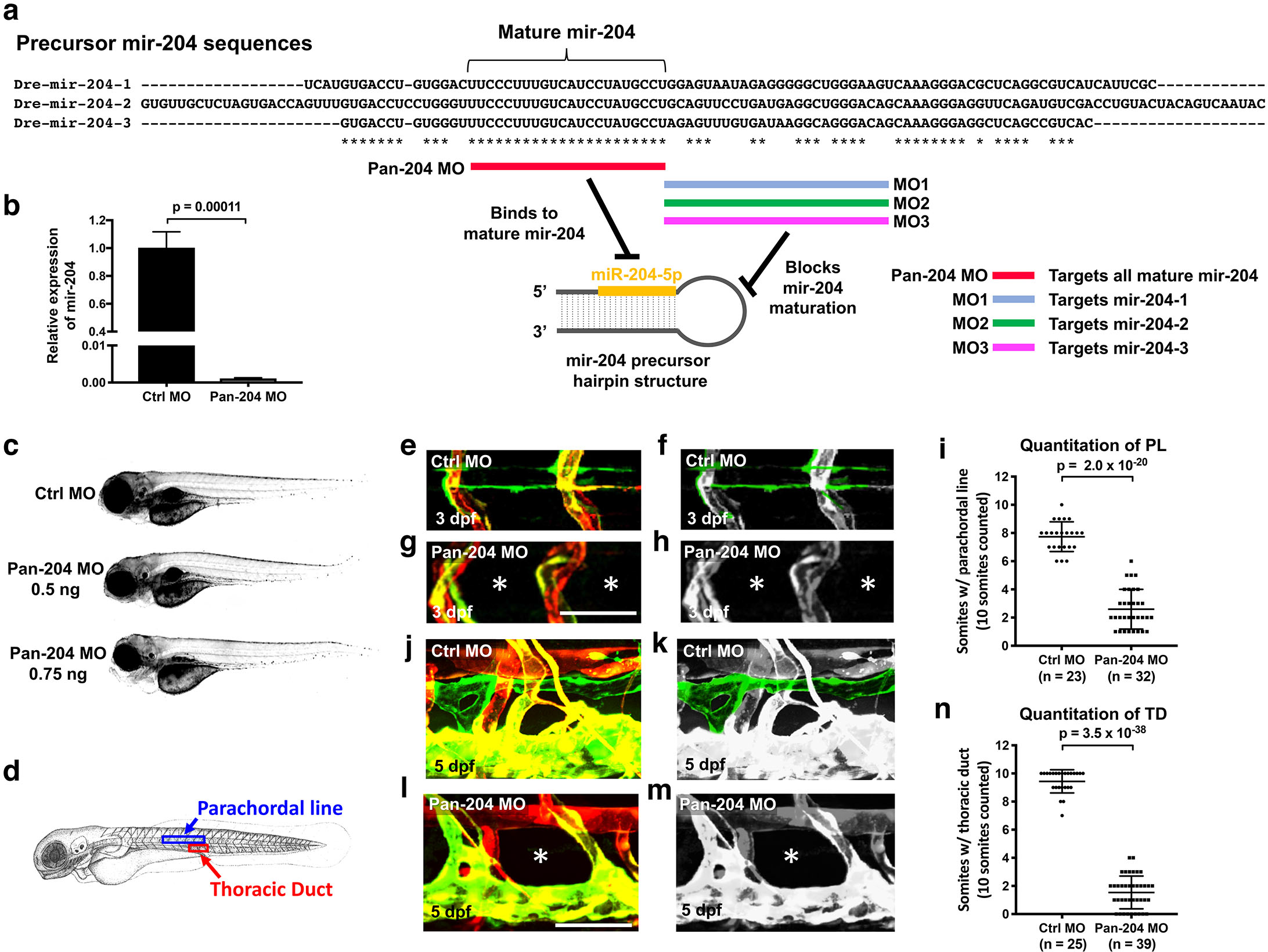
Defective lymphangiogenesis in mir-204 deficient zebrafish. **(a)**Sequence alignment of the three zebrafish precursor mir-204 sequences (*mir-204-1, mir-204-2,* and *mir-204-3*) and a schematic diagram showing four morpholinos (pan-204 MO, MO1, MO2, and MO3) targeting them. The data shown in the rest of this figure (panels b-m) uses the pan-204 MO targeting the mature mir-204 sequence generated by all three zebrafish mir-204 loci. **(b)**Quantitative TaqMan RT-PCR measurement of the relative levels of mature miR-204 in 1 dpf control MO- or pan-204 MO-injected embryos, normalized to controls. Three biological replicates were analyzed. **(c)**Bright field microscopic images of 5 dpf zebrafish larve that were injected with 0.5 ng of control MO (top) 0.5 ng of pan-204 MO (middle), or 0.75 ng of pan-204 MO (bottom). **(d)**Schematic of a zebrafish larva indicating the locations imaged in panels e-h (parachordal line) and j-m (thoracic duct). **(e-h)**Confocal images of the parachordal line in 3 dpf animals injected with either control MO (e, f) or pan-204 MO (g, h). In panels f and h the parachordal line is highlighted in green and other vessels are in grey. The absence of the parachordal line is noted with asterisks in panels g and h. **(i)**Quantification of parachordal line formation in 3 dpf animals injected with either control MO (n = 23) or pan-204 MO (n = 32). A total of 10 somitic segments were scored in each animal for the presence or absence of an intact parachordal line. **(j-m)**Confocal images of the thoracic duct in 5 dpf animals injected with either control MO (j, k) or pan-204 MO (l, m). In panels k and m the thoracic duct is highlighted in green and other vessels are in grey. The absence of the thoracic duct is noted with asterisks in panels l and m. **(n)**Quantification of thoracic duct formation in 5 dpf animals injected with either control MO (n = 25) or pan-204 MO (n = 39). A total of 10 somitic segments were scored in each animal for the presence or absence of an intact thoracic duct. All images are lateral views of MO-injected *Tg(mrc1a:eGFP)*^*y251*^, *Tg(kdrl:mCherry)*^*y171*^ double-transgenic animals. Scale bar: 50 μm (g, l). All graphs are analyzed by t-test and the mean ± SD is shown.

### mir-204 function is required for lymphatic development

To further examine the role of mir-204 in lymphatic development we generated a mutant in *mir-204-1* (the conserved mir-204 located in intron 5 of *trpm3;* **Figure 2-figure supplement 1a**) via CRISPR/Cas9 technology. We identified an active single guide RNA (sgRNA) targeting *mir-204-1* (**Figure 2**), and used this sgRNA to generate a 22 bp deletion including the majority of the mature mir-204 sequence (17 out of 22 nt) and the entire seed sequence of mir-204 (**Figure 3b**). However, *mir-204-1* homozygous mutants (*miR-204-1*^*−/−*^) showed only an approximately 20% decrease in mir-204 levels compared to wildtype embryos, supporting the idea that reduction in *mir-204-1* can be compensated for by *mir-204-2* and/or *mir-204-3* (**Figure 3c**). As might be expected from this modest reduction, *mir-204-1*^*−/−*^ animals were viable and fertile and did not have obvious morphological defects up to adulthood. However, injecting *mir-204-1*^*−/−*^ mutants with MO2 and MO3 (targeting *mir-204-2* and *mir-204-*3, respectively; **Figure 3d**) resulted in a 90% reduction in mir-204 levels compared to wildtype animals (**Figure 3e**). Although *mir-204-1*^*−/−*^ mutants or wild type animals co-injected with MO2 and MO3 form normal thoracic ducts similar to those in uninjected wild type animals, *mir-204-1*^*−/−*^ mutant animals co-injected with MO2 and MO3 fail to properly form the thoracic duct and other lymphatic vessels (**Figure 3f-j**), like pan-204 MO-injected animals (**Figure 2**).

**Figure 3.**
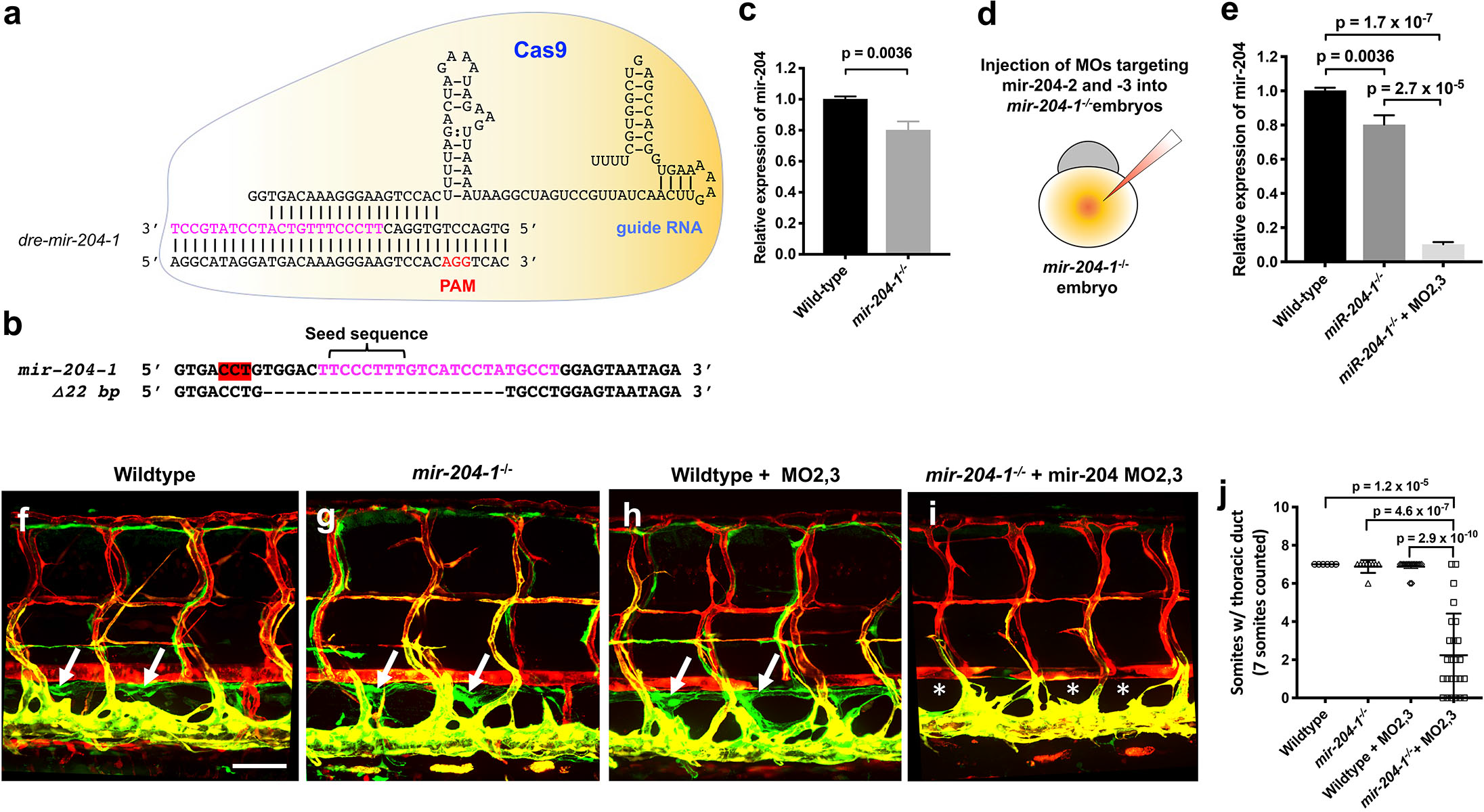
Mir-204 function is required for lymphatic development. **(a)**Schematic of CRISPR/Cas9 and guide RNA targeting of *mir-204-1*. **(b)**Sequence alignment of wildtype and *mir-204-1* mutant genomic DNA. The mature mir-204 sequence is noted in magenta, and the PAM sequence is highlighted in red. The mutant carries 22 bp deletion that removes 17 nucleotides of the mature mir-204 sequence. **(c)**Quantitative TaqMan RT-PCR measurement of the relative levels of mature mir-204 in 1 dpf *mir-204-1*^*−/−*^ mutant and wild type sibling embryos, normalized to wild type levels. Three biological replicates were analyzed. **(d)**Schematic diagram illustrating MO2 and MO3 co-injection into a 1-cell stage *mir-204-1*^*−/−*^ mutant embryo. **(e)**Quantitative TaqMan RT-PCR measurement of the relative levels of mature mir-204 in 1 dpf wild type sibling, *mir-204-1*^*−/−*^ mutant, and MO2 + MO3 co-injected *mir-204-1*^*−/−*^ mutant animals. **(f-i)**Representative confocal images of the mid-trunk of 5 dpf wild type sibling (f), *mir-204-1*^*−/−*^ mutant (g), MO2 + MO3 co-injected wild type sibling (h), and MO2 + MO3 co-injected *mir-204-1*^*−/−*^ mutant (i) animals. Images are lateral views of *Tg(mrc1a:eGFP)*^*y251*^, *Tg(kdrl:mCherry)*^*y171*^ double-transgenic animals, rostral to the left. The thoracic duct is labelled with white arrows, and absence of the thoracic duct is noted with asterisks. **(j)**Quantification of thoracic duct formation in 5 dpf wild type (n = 6), *mir-204-1*^*−/−*^ mutant (n = 9), MO2 + MO3 co-injected wild type (n = 16), and MO2 + MO3 co-injected *mir-204-1*^*−/−*^ mutant animals (n = 25). The number of somitic segments with an intact thoracic duct was counted, with the same 7 mid-trunk somites measured in each animal. Scale bar: 100 μm (f). All graphs are analyzed by t-test and the mean ± SD is shown.

To further examine the relative importance of each of the zebrafish mir-204s, we carried out additional experiments using the MOs targeting the dicer-cleavage sites of each of the three zebrafish mir-204s (**Figure 2a and Figure 3-figure supplement 1**). The sequences targeted by these MOs (MO1, MO2, MO3) differ from one another by 27-46% (**Figure 3-figure supplement 1a**). Injection of 0.5 ng of each individual MO alone did not affect lymphatic vessel development (**Figure 3-figure supplement 1b-d,i**). Pairwise co-injection of 0.5 ng MO1 and 0.5 ng MO2 resulted in defective thoracic duct formation (**Figure 3-figure supplement 1e,j**), but pairwise co-injection of either MO1 + MO3 or MO2 + MO3 did not affect lymphatic vessel formation (**Figure 3-figure supplement 1f,g,j**). Animals injected with 0.5 ng of each of the three MOs (MO1, MO2, and MO3, for a total of 1.5 ng injected) had lymphatic defects similar to but no more severe than animals injected with either the MO1 + MO2 combination or 0.5 ng of pan-204 MO (**Figure 3-figure supplement 1h,j and Figure 2n**). These results suggest that *mir-204-1* and *mir-204-2* play relatively more important roles than *mir-204-3* during lymphatic development, and confirm that suppressing mir-204 function leads to defective lymphatic vessel formation.

### Endothelial expression of mir-204 drives precocious lymphangiogenesis

To further examine the role of mir-204, we used the mrc1a promoter (Jung et al., 2017) to drive mir-204 expression in venous and lymphatic endothelial cells using a previously reported transgene vector (Nicoli et al., 2010) containing splice donor (SD) and splice acceptor (SA) sequences from the EF1alpha gene, with the *dre-mir-204-1-*containing trpm3 intron 5 sequence cloned in between them (**Figure 4a**). We isolated germline transgenic animals carrying this *Tg(mrc1a:mir204-egfp)* transgene. Adult germline transgenic animals are viable and fertile with no obvious superficial morphological abnormalities, but germline transgenic progeny generated from these fish display precocious development of the thoracic duct (**Figure 4b-d**). Normal *Tg(mrc1a:eGFP)* animals begin to form the thoracic duct at approximately 3-4 dpf, with a nearly complete thoracic duct formed by around 4-5 dpf (**Figure 4b**). However, in *Tg(mrc1a:mir204-eGFP)* animals the thoracic duct forms precociously, with a nearly complete thoracic duct already present by 3 dpf (**Figure 4c,d**), suggesting that increased mir-204 expression is promoting early lymphatic development.

**Figure 4.**
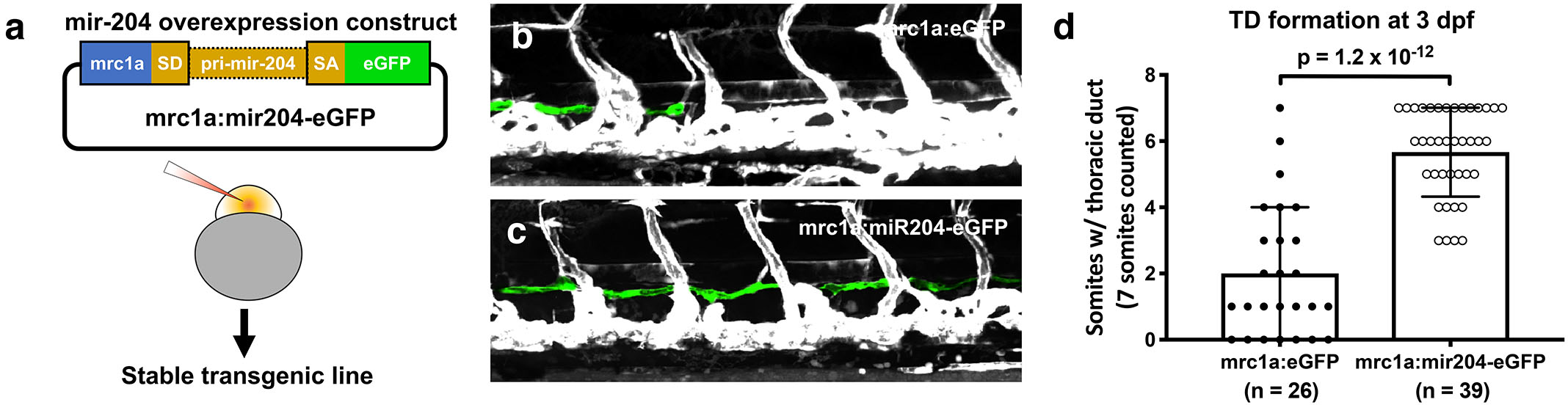
Endothelial expression of mir-204 drives precocious lymphatic development. **(a)**Schematic illustration of the mir-204 expression construct. The EGFP expression cassette is driven by the *mrc1a* promoter (Jung et al., 2017), with ~1kb of *dre-miR-204-1* genomic sequence from the fifth intron of the *trpm3* gene cloned between splice donor (SD) and splice acceptor (SA) sequences and flanking exonic sequences from the *ef1a* gene. The original vector backbone was previously described (Nicoli et al., 2010). Stable transgenic lines were generated by injecting this construct into wild type animals. **(b,c)**Representative confocal images of mid-trunk of 3 dpf *Tg(mrc1a:eGFP)* control (b) and *Tg(mrc1a:mir204-eGFP)* mir-204-expressing (c) germline transgenic animals. The thoracic duct is pseudocolored in green, with other vessels in grey. **(d)**Quantification of thoracic duct formation in 3 dpf *Tg(mrc1a:eGFP)* control and *Tg(mrc1a:mir204-eGFP)* mir-204-expressing germline transgenic animals. The number of somitic segments with an intact thoracic duct was counted, with the same 7 mid-trunk somites measured in each animal. All graphs are analyzed by t-test and the mean ± SD is shown.

### NFATC1 is a conserved target of miR-204

Nuclear Factor of Activated T Cells 1 (NFATC1) is expressed in developing lymphatic endothelial cells and genetic ablation of Nfatc1 in mice causes abnormal lymphatic vessel patterning and lymphatic hyperplasia (Norrmen et al., 2009). We identified human NFATC1 as a potential miR-204 target with a single putative miR-204 binding site in its 3’UTR using RNA22 (Miranda et al., 2006) (**Figure 5a**). The NFATC1 3’UTR was cloned downstream of a luciferase reporter and this plasmid DNA construct was co-transfected into HEK293 cells together with either miR-204 or (as a negative control) miR-126. Luciferase reporter activity was significantly suppressed in the presence of miR-204 but not in the presence of the miR-126 control (**Figure 5b**). To confirm the specificity of NFATC1 miR-204 target site recognition, we mutated 4 nucleotides in the seed binding sequence of the NFATC1 3’UTR (**Figure 5a**) and demonstrated that this rendered the construct insensitive to suppression by miR-204 (**Figure 5b**), confirming that this binding site is critical for direct targeting. Overexpression of miR-204 in human LECs suppressed endogenous NFATC1 expression while miR-204 inhibitor (antagomir) increased endogenous NFATC1 expression, showing that miR-204 also regulates endogenous NFATC1 transcript levels in human LEC (**Figure 5c**). We observed similar regulation of *nfatc1* by mir-204 in the zebrafish. Using RNA22, we also found that zebrafish *nfatc1* has a single mir-204 binding site in its 3’UTR (**Figure 5d**). The activity of a luciferase reporter containing the zebrafish *nfatc1* 3’UTR is also strongly suppressed by miR-204, and this suppression is fully rescued by mutating the seed binding sequence in the *nfatc1* 3’UTR (**Figure 5e**). Importantly, mir-204 knockdown using the pan-204 MO (targeting all three zebrafish mir-204s) results in increased levels of endogenous *nfatc1* transcript in 5 dpf zebrafish embryos (**Figure 5f**). Together, these data demonstrate conserved miR-204-mediated regulation of NFATC1.

**Figure 5.**
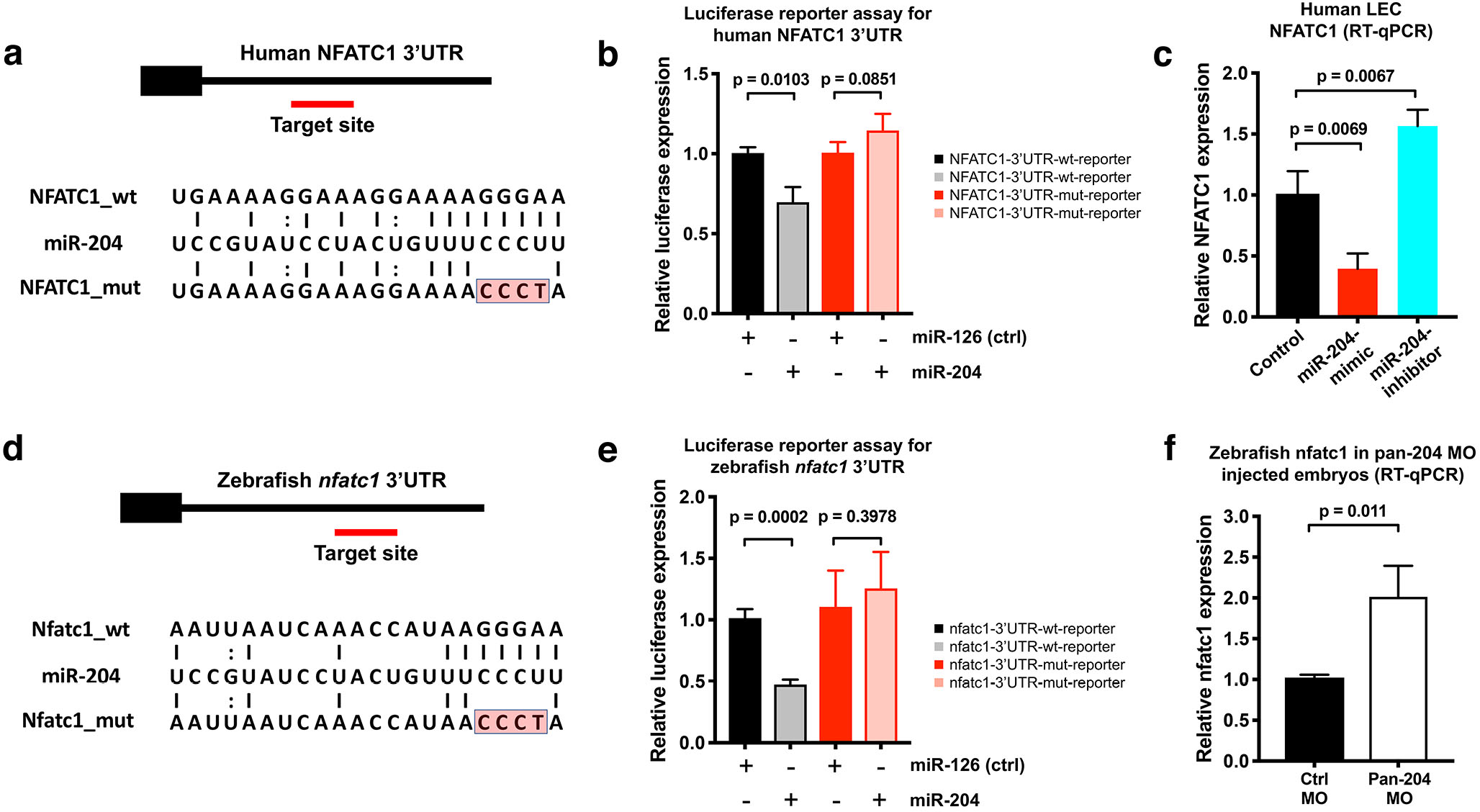
NFATC1 is a conserved target of miR-204. **(a)**Sequence alignment of mature miR-204 (middle line) and its target region in the human NFATC1 3’UTR (top line). A mutant form of the human NFATC1 3’UTR used for the luciferase assay in panel b is also shown (bottom line; 4 mismatches in the seed binding region are highlighted in red). **(b)**Quantitative luciferase reporter assay using wild type or mutant forms of the human NFATC1 3’UTR transfected into HEK293 cells together with either miR-204 or miR-126 (control). Four biological replicates were analyzed. **(c)**Quantitative TaqMan RT-PCR measurement of relative endogenous NFATC1 transcript levels in human LEC (HMVEC-dLy) transfected with miR-204-mimic or miR-204-antagomir, normalized to control mock transfected levels. **(d)**Sequence alignment of mature miR-204 (middle line) and its target region in the zebrafish nfatc1 3’UTR (top line). A mutant form of the zebrafish nfatc1 3’UTR used for the luciferase assay in panel e is also shown (bottom line; 4 mismatches in the seed binding region are highlighted in red). **(e)**Quantitative luciferase reporter assay using wildtype or mutant forms of the zebrafish nfatc1 3’UTR co-transfected into HEK293 cells together with either miR-204 or miR-126 (control). Four biological replicates were analyzed. **(f)**Quantitative TaqMan RT-PCR measurement of relative endogenous zebrafish nfatc1 transcript levels in 5 dpf animals that were injected with either control MO or pan-204 MO. Three biological replicates were analyzed. All graphs are analyzed by t-test and the mean ± SD is shown.

### Suppression of NFATC1 causes thoracic duct hyperplasia in the zebrafish

As noted above, loss of Nfatc1 in mice is associated with lymphatic hyperplasia (Norrmen et al., 2009). Using morpholinos targeting *nfatc1*, we were able to show that knockdown of this gene in the zebrafish also causes thoracic duct enlargement (**Figure 6a-e**). The calcineurin inhibitor cyclosporin A (CsA) blocks signaling downstream from NFAT, and treatment of mice with CsA during embryonic stages phenocopies the lymphatic effects caused by genetic ablation of Nfatc1 (Norrmen et al., 2009). By treating developing zebrafish larvae with a low dose of CsA (1 ug/mL), we were also able to phenocopy the thoracic duct enlargement phenotype observed in nfatc1 MO-injected animals (**Figure 6f-j**). Therefore, consistent with previous data in mice, our results suggest that nfatc1 is required for proper lymphatic development in the zebrafish.

**Figure 6.**
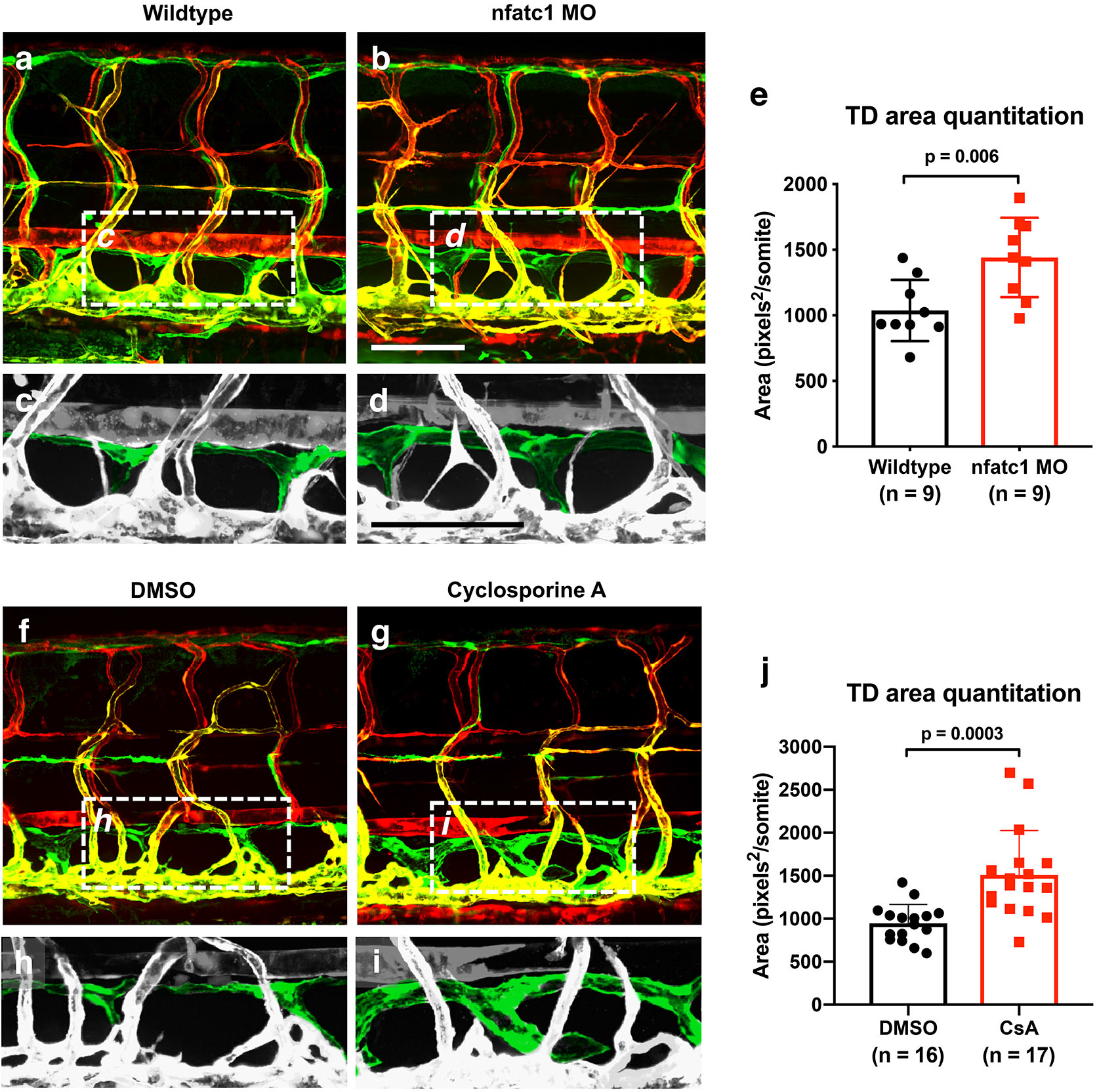
Suppression of Nfatc1 causes enlargement of the thoracic duct. **(a,b)** Confocal images of the mid-trunk of 5 dpf wildtype (a) and nfatc1 MO-injected (b) animals. The dashed boxes in panels a and b show the areas magnified in panels c and d, respectively. **(c,d)**Magnified images from panels a and b, with the thoracic duct pseudocolored in green and other vessels in grey. **(e)**Quantitation of thoracic duct size measured as the area encompassed by the thoracic duct in confocal images of the same 7 mid-trunk somitic segments in 5 dpf wildtype (n = 9) and nfatc1 MO-injected (n = 9) animals. **(f,g)**Confocal images of the mid-trunk of 5 dpf DMSO treated (f) and cyclosporine A (CsA) treated (g) animals. The dashed boxes in panels f and g show the areas magnified in panels h and i, respectively. **(h,i)**Magnified images from panels f and g, with the thoracic duct pseudocolored in green and other vessels in grey. **j,** Quantitation of thoracic duct size measured as the area encompassed by the thoracic duct in confocal images of 7 somitic segments from 5 dpf DMSO-treated (n = 16) and CsA-treated (n = 17) animals. All images are lateral views of *Tg(mrc1a:eGFP)*^*y251*^, *Tg(kdrl:mCherry)*^*y171*^ double-transgenic animals, rostral to the left. Scale bar = 100 μm (b,d). All graphs are analyzed by t-test and the mean ± SD is shown.

### Suppression of nfatc1 rescues lymphatic development in mir-204-deficient zebrafish

The results above suggested that increased expression of nfatc1 could be at least partially responsible for the defects in lymphatic vessel development in mir-204-deficient animals. To test this idea, we examined whether lymphatic vessel formation in mir-204-deficient animals could be “rescued” by nfatc1 knockdown. As shown above, pan-204 MO-injected animals fail to form the thoracic duct (**Figure 7a,b,d,e,j**) and dorsal longitudinal lymphatic vessel (DLLV) in the trunk (**Figure 7a,b,g,h,k**). However, in animals co-injected with both pan-204 MO and nfatc1 MO formation of the thoracic duct is largely restored, as well as other lymphatic vessels including the DLLV and intersegmental lymphatics (ISLV) (**Figure 7c,f,I,j,k**). These results suggest that abnormal lymphatic vessel development in mir-204 deficient animals can be substantially rescued by suppressing nfatc1 expression. Based on all of our findings, we propose that a proper balance between mir-204 and nfatc1 is critical for proper lymphatic vessel development, with loss of either mir-204 or nfatc1 causing defects in lymphatic vessel formation (**Figure 7l**).

**Figure 7.**
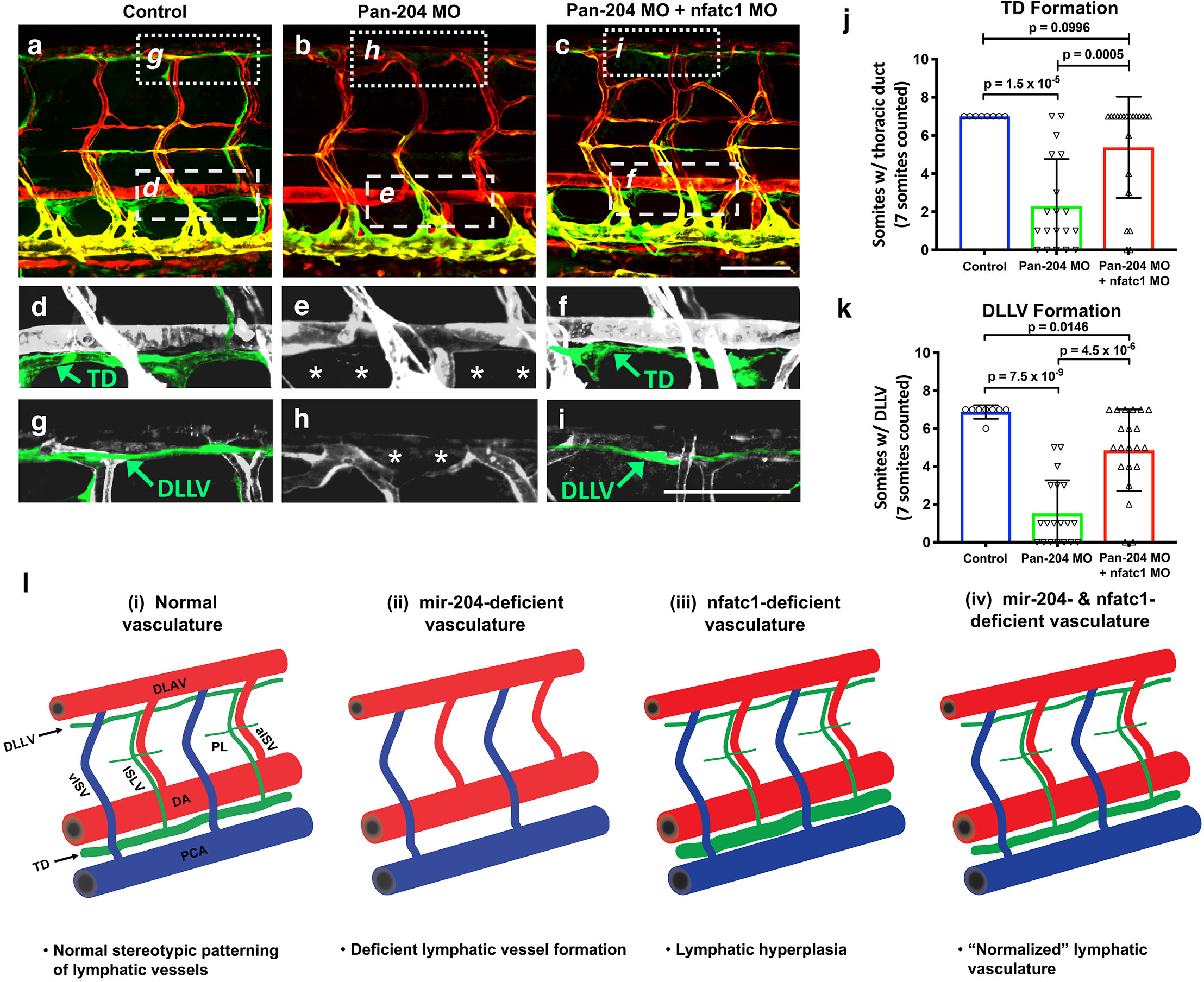
Suppression of nfatc1 rescues the lymphatic defects in mir-204-deficient animals. **(a-c)**Confocal images of the mid-trunk of 5 dpf control (a) pan-204 MO-injected (b) or pan-204 MO and nfatc1 MO co-injected (c) animals. White dotted boxes in panels a-c show areas magnified in panels d-f, respectively, while white dashed boxes show areas magnified in panels g-i, respectively. **(d-f)**Magnified images from panels a-c with the thoracic duct (TD) pseudocolored in green and other vessels in grey. The TD is labeled, and the absence of the TD is noted with asterisks. **(g-i)**Magnified images from panels a-c with the dorsal longitudinal lymphatic vessel (DLLV) pseudocolored in green and other vessels in grey. The DLLV is labeled, and the absence of the DLLV is noted with asterisks. **(j)**Quantification of thoracic duct (TD) formation in 5 dpf control (n = 8), pan-204 MO-injected (n = 19), or pan-204 MO and nfatc1 MO co-injected animals (n = 21). A total of 7 mid-trunk somitic segments were scored in each animal for the presence or absence of an intact TD. **(k)**Quantification of dorsal longitudinal lymphatic vessel (DLLV) formation in 5 dpf control (n = 8), pan-204 MO-injected (n = 19), or pan-204 MO and nfatc1 MO co-injected animals (n = 21). A total of 7 mid-trunk somitic segments were scored in each animal for the presence or absence of an intact DLLV. **(l)**Schematic diagrams illustrating 5 dpf zebrafish trunk lymphatic vessels present in (i) normal control, (ii) mir-204 deficient, (iii) nfatc1-deficient, and (iv) mir-204- and nfatc1-deficient animals. Suppression of mir-204 leads to loss of lymphatic vessels (ii), while nfatc1 deficiency causes lymphatic (thoracic duct) hyperplasia (iii). The lymphatic defects in mir-204 deficient animals can be rescued by simultaneous suppression of nfatc1 (iv). All images are lateral views of *Tg(mrc1a:eGFP)*^*y251*^, *Tg(kdrl:mCherry)*^*y171*^ double-transgenic animals, rostral to the left. Scale bar = 100 μm (c,i). All graphs are analyzed by t-test and the mean ± SD is shown.

## DISCUSSION

During early embryonic development Sox18 and Prox1 promote early lymphatic specification in a subset of primitive venous endothelial cells. These lymphatic endothelial progenitors turn on Vegfr3 expression and then migrate away from the veins to form primitive lymphatic structures (Yang and Oliver, 2014). Over the past few decades, characterization of the transcriptional programming directing lymphatic vessel formation has provided insights into molecular pathways regulating lymphangiogenesis during early development. However, the post-transcriptional steps that help to refine this tightly regulated process remain largely unexplored.

MicroRNAs (miRNAs) provide genetic robustness by fine-tuning and buffering gene expression to maximize gene regulation accuracy (Bartel, 2009, Schmiedel et al., 2015, Cassidy et al., 2013, Herranz and Cohen, 2010, Ebert and Sharp, 2012). Although previous studies have identified tissue-specific miRNAs, uncovering specific *in vivo* functions for miRNAs has been challenging, with modest or absent phenotypes for many miRNA-deficient animal models (Lai, 2015). In some cases, miRNA mutants have been shown to exert minor influences on organ development that are exacerbated under certain environmental stress condition. For example, animals lacking members of the miR-139 or miR-24 endothelial miRNA families display mild or no phenotype, but they do display greater phenotypic variability upon angiogenic stress, high temperature, or hypoxic conditions, suggesting these endothelial miRNAs help limiting phenotypic variation in the vascular system (Kasper et al., 2017). A recent systematic study of *Drosophila* miRNA knockouts reported that 80% of miRNA mutants display at least one observable defect throughout the course of development from embryogenesis to adulthood, indicating that despite their relatively modest phenotypes and variable degree of severity most *Drosophila* miRNA mutants do display some abnormalities (Chen et al., 2014). Recently, there have been efforts to identify endothelial miRNAs required for proper vascular network formation during vertebrate development. The best-characterized of these is probably the pan-endothelial miRNA miR-126. Loss of miR-126 results in leaky vessels and hemorrhage due to loss of vascular integrity and defects in endothelial function caused by misregulation of molecules in the VEGF pathway (Fish et al., 2008, Wang et al., 2008, Nicoli et al., 2010, Sessa et al., 2012). Although different groups have reported slightly different specific phenotypes (e.g. edema, hemorrhage, or embryonic lethality) (Wang et al., 2008, Fish et al., 2008, Kontarakis et al., 2018, Chen et al., 2016), a recent loss-of-function study using mice and zebrafish showed that miR-126 also plays a role in lymphangiogenesis by modulating flt4 signal transduction (Kontarakis et al., 2018).

To identify candidate LEC-enriched miRNAs with potentially more specific lymphatic functions, we used small RNA profiling of human primary endothelial cells. For LECs we used HMVEC-dLy, while for BECs we used HUVEC. Both of these cells have been previously well-validated as being representative of BEC and LEC, respectively (Pedrioli et al., 2010, Dunworth et al., 2014), and we confirmed that they have appropriate differential expression of well-characterized markers of blood and lymphatic endothelial identity. Using small RNA sequencing, we identified 30 highly significantly LEC-enriched miRNAs. Interestingly, the majority of these miRNAs were specific to mammals. Only 7 miRNAs were conserved in other vertebrate species, suggesting that most of these LEC miRNAs may have diverged and/or evolved for mammalian-specific functions. However, the most LEC-enriched miRNA was miR-204, a highly conserved miRNA across a broad swath of vertebrate species, suggesting it plays an important conserved role during lymphatic development. In addition to miR-204 and a number of other newly identified LEC-enriched miRNAs, we also uncovered miRNAs such as miR-326, miR-139, miR-338, miR-148a, and miR-30d that had been previously reported to be LEC-enriched (Pedrioli et al., 2010). Intriguingly, miR-204 was not identified in the previous study by Pedrioli et al, despite its being the most highly enriched lymphatic microRNA in our study, while the most enriched LEC miRNA reported by Pedrioli et al, miR-95, was excluded from our analysis due to a low number of sequence reads. The reasons for the disparate results obtained from our data set and that of Pedrioli et al. are not clear, but they may reflect differences in the starting material for sequencing, the profiling methods used, or the bioinformatic tools and filters employed. We would note that out study used three biological replicates for both the LEC and BEC sequencing, and that we sequenced to a relatively high depth and applied very stringent filtering to our data set.

Our study establishes a critical role for miR-204 during lymphatic development. As noted above, mature miR-204 displays a remarkable evolutionary conservation throughout the vertebrates, with 100% sequence identity between mammals, avians, and fish, suggesting that this miRNA likely plays an important conserved function. Indeed, our zebrafish results show that suppression of mir-204 leads to strong defects in lymphangiogenesis. Although the sequence of mature zebrafish mir-204 is identical to human miR-204, human miR-204 is encoded by only a single locus in intron 6 of the TRPM3 gene, while zebrafish mir-204 is encoded by three separate loci within the introns of the *trpm3, trpm1a,* and *trpm1b* genes. Interestingly, the mammalian *TRPM1* gene has a related miRNA encoded within a similar intron as in zebrafish *trpm1a,* and *trpm1b* that may have evolved from mir-204. We used several complementary mir-204 loss-of-function approaches in the zebrafish, including (i) a morpholino targeting the mature mir-204 sequence produced by all three zebrafish mir-204’s, (ii) a CRISPR-Cas9-generated genetic mutant ablating *mir-204-1*, and (iii) morpholinos individually targeting sequences required for the maturation of each of the three zebrafish mir-204’s, which were co-injected with one another or with the *mir-204-1* mutant. Our results revealed that all treatments that target at minimum *mir-204-1* and *mir-204-2* result in comparable dramatic loss-of-lymphatic phenotypes. The cells forming the parachordal line at 2-3 dpf are the earliest lymphatic endothelial cells to emerge in the zebrafish, and mir-204 suppression results in loss of the parachordal line. By 5 dpf lymphatic progenitors derived from the parachordal line form the thoracic duct, the major vertebrate trunk lymphatic vessel, and thoracic duct formation is also largely absent in mir-204 deficient animals.

Our study also identified a key downstream target of miR-204 regulation during lymphangiogenesis. Previous studies have shown that the Nuclear Factor of Activated T-cells (NFAT) protein family member NFATC1 is important for lymphatic vessel development (Norrmen et al., 2009, Kulkarni et al., 2009). NFATC1 is co-expressed with Prox1-, Vegfr3-, and Pdpn-positive LEC progenitors originating from the cardinal vein indicating their potential involvement in lymphatic specification, and Nfatc1 null mice develop lymphatic hyperplasia suggesting a role in lymphatic maturation (Norrmen et al., 2009, Kulkarni et al., 2009). Consistent with these previous data, we showed that nfatc1 knockdown results in similar lymphatic enlargement in the zebrafish. We identified NFATC1 as a probable miR-204 target based on conserved miR-204 binding sites in the 3’ UTRs of both human and zebrafish NFATC1 transcripts and the expression of NFATC1 was suppressed by miR-204 mimic and increased by miR-204 antagomir in human LECs. We were able to verify targeting of the NFATC1 3’ UTR by miR-204 using firefly/renilla dual luciferase NFATC1 3’ UTR reporter assays, and demonstrate up-regulation of zebrafish nfatc1 upon mir-204 knockdown, confirming that miR-204 suppresses NFATC1 levels. Finally, we showed that knocking down nfatc1 could rescue lymphatic development in mir-204-deficient zebrafish. These results suggest that balanced expression of mir-204 and nfatc1 is critical for proper developmental lymphangiogenesis.

Since miRNAs often regulate many targets, and miR-204 does have potential target binding sites in other genes, further investigation will be required to characterize some of these additional genes and determine whether some of the lymphatic effects of miR-204 are mediated via regulation of other targets in addition to nfatc1. Nevertheless, since NFATC1 has already been shown to play an important role in lymphangiogenesis in mammals, and since suppression of nfatc1 in the zebrafish causes lymphatic hyperplasia, and can effectively rescue the effects of miR-204 knockdown, our results suggest that nfatc1 is a major, key downstream target regulated by miR-204 in developing lymphatic endothelial cells.

In summary, our work establishes an important role for miR-204 in regulating lymphatic vascular network formation during embryonic development via modulation of its conserved target NFATC1. Our findings provide important new insight into the role of lymphatic-enriched miRNAs during developmental lymphangiogenesis, and by unveiling these pathways provide new opportunities to understand lymphatic development and associated disorders.

## MATERIALS AND METHODS

### Zebrafish and drug treatment

Zebrafish were maintained and zebrafish experiments were performed according to standard protocols (Westerfield, 2000) and in conformity with the Guide for the Care and Use of Laboratory Animals of the National Institutes of Health, in an Association for Assessment and Accreditation of Laboratory Animal Care (AAALAC) accredited facility. The *Tg(mrc1a:eGFP)*^*y251*^;*Tg(kdrl:mCherry)*^*y171*^ double transgenic line was used in this study (Jung et al., 2017). For nfatc1 inhibition, 24 hpf embryos were dechorinated and incubated in 1 ug/mL cyclosporine A (CsA) or DMSO for 4 days and the animals imaged at 5 dpf.

### Transgenic constructs and animals

The Tol2(mrc1a:mir204-eGFP) venous/lymphatic endothelial autonomous expression construct was generated using Tol2kit components with Gateway Technology (Kwan et al., 2007). To make a *mir-204* middle entry cassette for the overexpression construct, we used pME-miR, containing an partial EF1alpha gene exon 1, intron 1, and exon 2 (Nicoli et al., 2010). A 1-kb genomic DNA sequence from intron 5 of the *trpm3* gene harboring the mir-204 precursor was amplified by PCR and subcloned into the multiple cloning site located in EF1alpha intron 1 of pME-miR using EcoR1 and Kpn1, to generate pME-miR204. To generate the final venous/lymphatic endothelial autonomous mir-204 expression construct, we combined p5E-mrc1a (Jung et al., 2017), pME-mir204, p3E-EGFPpA (Kwan et al., 2007), and pDestTol2pA(Kwan et al., 2007). The DNA construct was microinjected into the blastomere of one-cell stage zebrafish embryos to generate transgenic insertions. Injected animals were raised to adult hood and their progeny were screened for germline transmission and expression of the transgene.

### Flow cytometry

All embryos subjected to FACS sorting were raised in E3 medium. 5 dpf *Tg(mrc1a:egfp)*^*y251*^, *Tg(kdrl:mCherry)*^*y171*^ double transgenic zebrafish were anesthetized with MS-222 and washed with 1X PBS (pH 7.4, without Ca^2+^ and Mg^2+^) three times. Animals were deyolked by gentle pipetting in yolk dissociation solution (55mM NaCl, 1.8mM KCl, 1.25mM NaHCO3). Cells were then dissociated by gentle pipetting in 0.25% trypsin-EDTA and 50mg/mL collagenase solution. Dissociated cells were passed though 70μm filter and centrifuged at 4000 rpm for 5 min at room temperature. Cells were washed and resuspended with 1X PBS. Fluorescent cell sorting was performed on a BD FACS ARIA (Becton Dickinson, Franklin Lakes, NJ). Isolated GFP+, mCherry+, and double positive cells were pelleted at 2,500 rpm for 5 min.

### Morpholino injections

All morpholinos (MOs) used in this study were acquired from Gene Tools to specifically target mature mir-204 or precursor mir-204s. The nfatc1 MO sequence was obtained from a previous study (Tijssen et al., 2011). MOs were injected into one cell stage *Tg(mrc1a:egfp)*^*y251*^, *Tg(kdrl:mCherry)*^*y171*^ double transgenic zebrafish embryos (Jung et al., 2017). Injected embryos were allowed to develop at 28.5 ºC until being imaged at the desired stage. Morpholino doses used were determined by performing dose curves to establish the optimal dose to minimize off-target effects. All morpholino sequences are listed (**Supplementary file 1a**).

### Genome editing

CRISPR genome editing technology was used to generate mutants in *dre-miR-204-1.* Codon-optimized Cas9 plasmid pT3TS-nls-zCas9-nls was used as template to *in vitro* transcribe Cas9 mRNA (Jao et al., 2013). A single guide RNA (sgRNA) was designed to target within the precursor *mir-204-1* sequence using CRISPRscan (http://www.crisprscan.org) (Moreno-Mateos et al., 2015). Fluorescence PCR was performed using AmpliTaq Gold DNA polymerase (Life Technologies) with M13F primer with fluorescence tag (6-FAM), amplicon-specific forward primer with M13 forward tail (5’- TGTAAACGACGGCCAGT-3’) and 5’PIG-tailed (5’-GTGTCTT-3’) amplicon-specific reverse primer for initial genotyping to identify G0 alleles that contain larger than ~10bp seed sequence deletions (Varshney et al., 2015). An ABI 3730 Genetic Analyzer Avant (Thermofisher) was used to analyze the PCR products. A founder was identified with a 22 bp deletion removing most of the mature mir-204-1 sequence. Adult F1 progeny of these founder fish were genotyped using conventional gel electrophoresis on a 2.5% agarose gel. All oligos used for genome editing are listed (**Supplementary file 1b**).

### Cell culture and transfection

HMVEC-dLy cells (Lonza) were cultured in EGM-2 MV BulletKit (Lonza, CC-3202) that contains hEGF, hydrocortisone, GA-1000, FBS, VEGF, hFGF-B, R^3^-IGF-1, and ascorbic acid. HUVECs (Lonza) were cultured in bovine hypothalamus extract, 0.01% Heparin and 20% FBS in M199 base media (Gibco) on 1mg/mL gelatin-coated tissue culture flasks. HEK293 cells were cultured in Advanced DMEM supplemented with 10% FBS and antibiotics. Transfection was performed using Lipofectamine 2000 (Invitrogen) and miRNA mimics (Life Technologies).

### RNA isolation, small RNA-seq, and TaqMan PCR

RNA isolation was performed using the mirVana kit (Life Technoloiges). A NanoDrop ND-100 spectrophotometer (Nanodrop Technology Inc.), Qubit 2.0 fluorometer (Life Technologies Inc.) and Agilent 2100 bioanalyzer (Aglient) were used to analyze RNA quantity and quality. Small RNA sequencing was performed by ACGT, Inc. Three biological samples were subjected to analysis. Briefly, libraries were prepared using the Illumina TruSeq Small RNA Sample Preparation Kit, then sequenced on a NextSeq 500 Illumina instrument, generating 50 bp single end reads. Data was analyzed using PartekFlow analysis software (Partek, Inc.). The sequence reads were trimmed to remove the following adapters: GTTCAGAGTTCTACAGTCCGACGATC (from the 5’ end) and TGGAATTCTCGGGTGCCAAGG (from the 3’ end). Then, the bases at the end of the sequences with quality less than 20 were removed. The remaining sequences were aligned to the human genome browser (hg38) and miRbase mature microRNA version 22 using Bowtie. The data was filtered for the counts smaller than 10 in 50% of samples, and used CPM (counts per million) for normalization. Differential expression was analyzed by Partek GSA algorithm. All supplies for TaqMan microRNA/gene assays were purchased from Life Technologies, and qPCR was performed using a CFX96 (BioRad). 18S rRNA (for human cells) and ef1a (for zebrafish cells) were used for internal controls for mRNAs, and U6 snRNA was used as an internal control for miRNAs.

### Luciferase reporter assay

The human NFATC1 3’UTR was PCR amplified from cDNA generated from human LEC RNA, and the zebrafish nfatc1 3’UTR from zebrafish embryo RNA. 3’UTR sequences were cloned downstream from the *renilla* luciferase gene using the XhoI and NotI sites in the siCHECK-2 vector (Promega, Madison, WI). This vector also contains a *firefly* luciferase gene driven by an independent protomer, which serves an internal control for the assay. Mutations in the miR-204 binding sites were generated using the QuikChange site-directed mutagenesis kit (Stratagene, La Jolla, CA) and the mutated sequences were confirmed by DNA sequencing. All the primers used for generating luciferase constructs are listed (**Supplementary file 1c**). HEK293 cells transfected with luciferase reporters and miRNA mimics were harvested after 24 h. A dual luciferase reporter assay system was used to determine luciferase levels (Promega, Madison, WI).

### Imaging methods

Embryos were anesthetized using 1x tricaine and mounted in 0.8-1.5% low melting point agarose dissolved in embryos media and mounted on a depression slide (Jung et al., 2016). Confocal fluorescence imaging was performed with a Nikon Yokogawa CSU-W1 spinning disk confocal microscope. The images were analyzed using ImageJ (Schindelin et al., 2012), Imaris 7.4 (Bitplane), and Photoshop (Adobe) software.

### Study Approval

Zebrafish husbandry and research protocols were reviewed and approved by the NICHD Animal Care and Use Committee at the National Institutes of Health. All animal studies were carried out according to NIH-approved protocols, in compliance with the *Guide for the Care and use of Laboratory Animals*.

## Author Contributions

HMJ, CH, AF, AD, DC, VP, LP performed experiments; HMJ, CH, AF, and BMW analyzed results and prepared the figures; HMJ and BMW designed the research and wrote the paper.

## Acknowledgements

The authors would like to thank members of the Weinstein laboratory for their critical comments on this manuscript. This work was supported by the intramural program of the *Eunice Kennedy Shriver* National Institute of Child Health and Human Development, National Institutes of Health (ZIA-HD008808 and ZIA-HD001011, to BMW).

## FIGURE SUPPLEMENT LEGENDS

**Figure 1-figure supplement 1.**
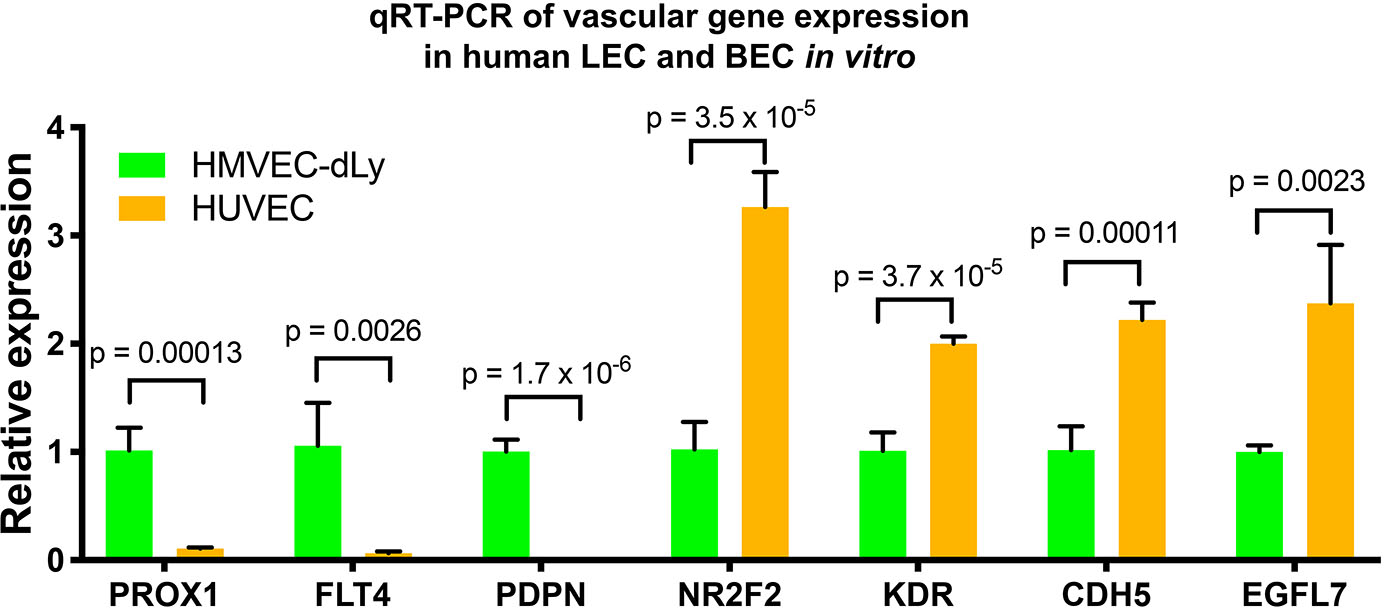
Differential expression of vascular genes in HMVEC-dLy and HUVEC. Quantitative TaqMan RT-PCR measurement of the relative expression of known lymphatic and blood vessel markers in HMVEC-dLy (Lymphatic Endothelial Cells, LEC) and HUVEC (Blood Endothelial Cells, BEC), normalized to expression levels in HMVEC-dLy (LEC). Four biological replicates were analyzed. All graphs are analyzed by t-test and the mean ± SD is shown.

**Figure 1-figure supplement 2.**
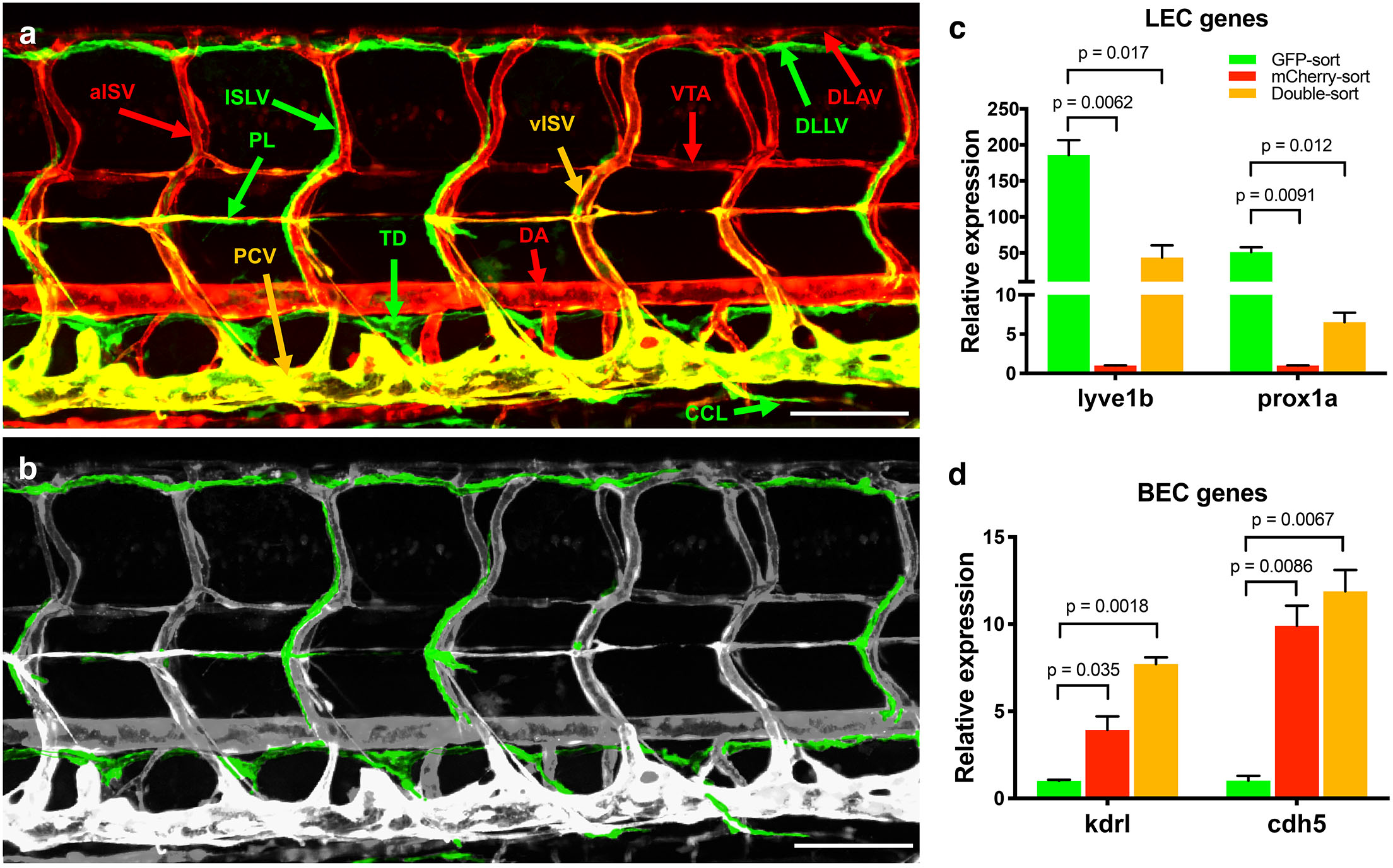
Trunk vascular patterning and vascular gene expression in FACS-sorted endothelial cells from transgenic zebrafish. **(a)**Confocal images of the vasculature in the mid-trunk of a 5 dpf *Tg(mrc1a:eGFP)*^*y251*^, *Tg(kdrl:mCherry)*^*y171*^ double-transgenic animal with mCherry positive arterial blood vessels (red), EGFP positive lymphatic vessels (green), and mCherry and EGFP double positive venous blood vessels (yellow). The different vessels are labeled: DA, dorsal aorta; DLLV, dorsal longitudinal lymphatic vessel; DLAV, dorsal longitudinal anastomotic vessel; ISLV: intersegmental lymphatic vessel; aISV: arterial intersegmental vessel; vISV: venous intersegmental vessel; PCV, posterior cardinal vein; PL, parachordal line; TD, thoracic duct; CCL, collateral cardinal lymphatics; VTA, vertebral artery. **(b)**Confocal image from panel A with lymphatic vessels pseudocolored in green and other vessels in grey, for easier visualization of the lymphatic network. **(c)**Quantitative TaqMan RT-PCR measurement of the relative expression of lymphatic endothelial cell (LEC) genes *lyve1b* and *prox1a* in FACS-sorted zebrafish endothelial cell populations. Expression is normalized to the arterial (mCherry-positive) endothelial cell population. **(d)**Quantitative TaqMan RT-PCR measurement of the relative expression of blood endothelial cell (BEC) genes *kdrl* and *cdh5* in FACS-sorted zebrafish endothelial cell populations. Expression is normalized to the lymphatic (EGFP-positive) endothelial cell population. Scale bars = 100 μm (a,b). Biological duplicates were analyzed. All graphs are analyzed by t-test and the mean ± SD is shown.

**Figure 2-figure supplement 1.**
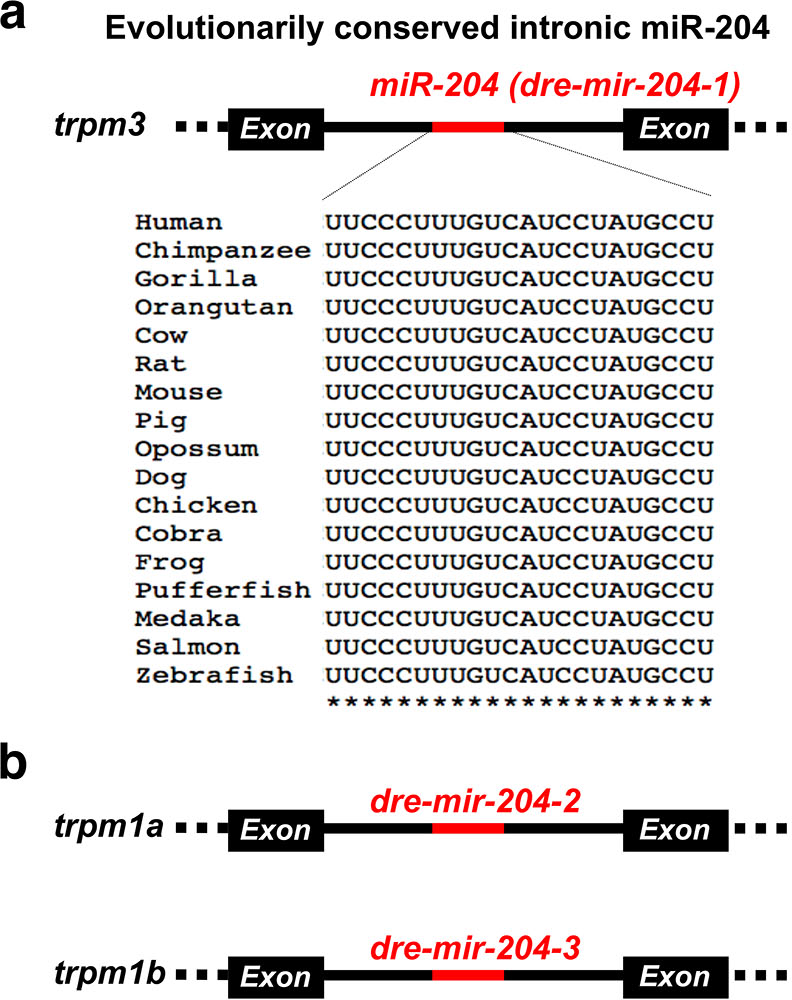
Evolutionarily conservation of miR-204. **(a)**Schematic diagram showing the location of miR-204 in intronic region of TRPM3 gene and the 100% conservation of its mature miRNA sequence amongst various vertebrate species. **(b)**Schematic of two additional zebrafish mir-204 loci located in intron 5 and 4 of *trpm1a* and *trpm1b,* respectively.

**Figure 3-figure supplement 1.**
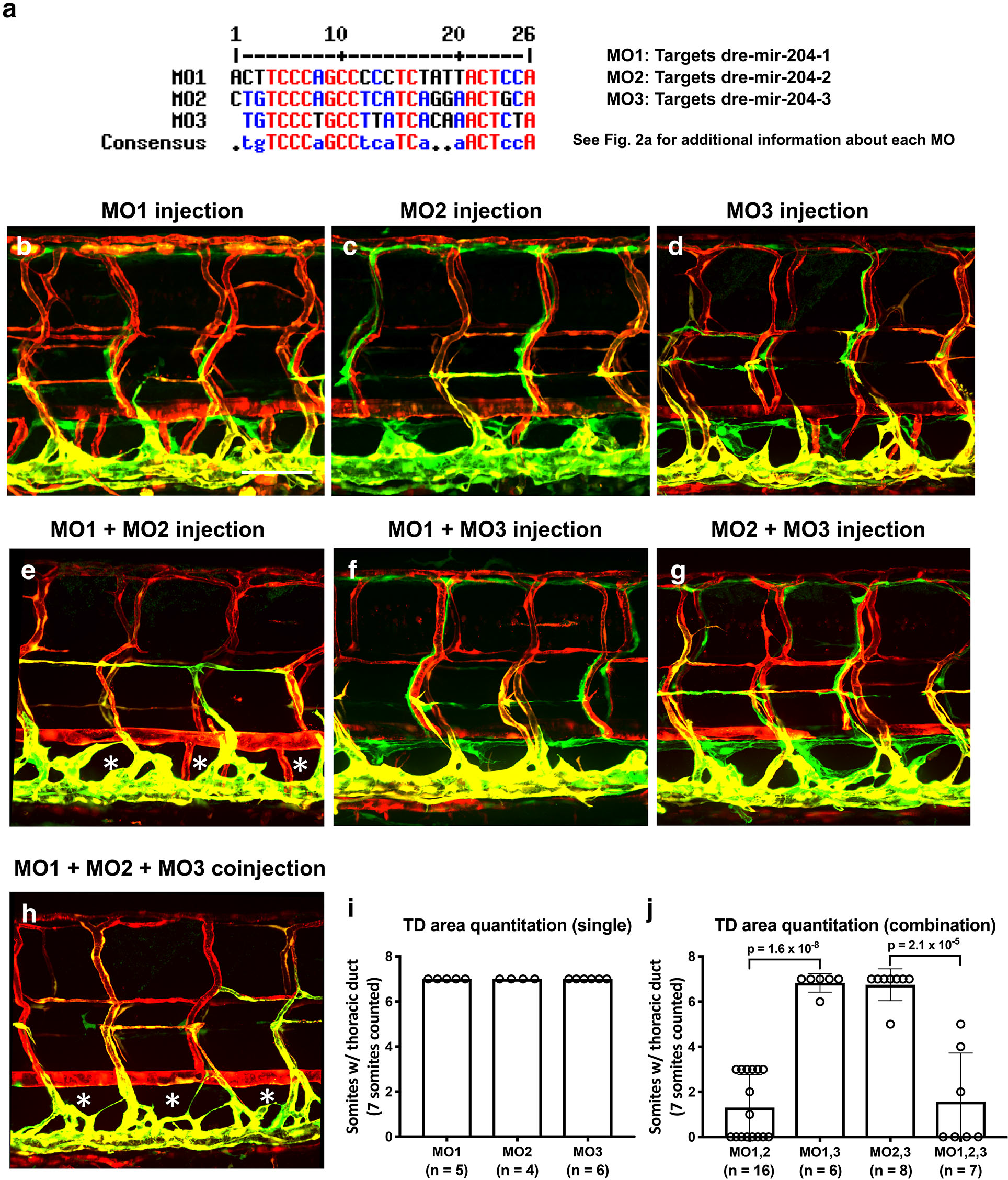
The effects of single or combined injections of mir-204 targeting MOs. **(a)**Sequence alignment of MO1, 2, 3. **(b-d)**Representative confocal images of 5 dpf *Tg(mrc1a:eGFP)*^*y251*^, *Tg(kdrl:mCherry)*^*y171*^ double-transgenic animals injected with 0.5 ng of MO1 (b), MO2 (c), or MO3 (d). **(e-g)**Representative confocal images of 5 dpf *Tg(mrc1a:eGFP)*^*y251*^, *Tg(kdrl:mCherry)*^*y171*^ double-transgenic animals injected pairwise with 0.5 ng each of MO1+MO2 (e), MO1+MO3 (f), or MO2+MO3 (g), for a final combined MO dose of 1.0 ng in each case. **(h)**Representative confocal image of a 5 dpf *Tg(mrc1a:eGFP)*^*y251*^, *Tg(kdrl:mCherry)*^*y171*^ double-transgenic animal injected with 0.5 ng each of MO1+MO2+MO3 (for a final combined MO dose of 1.5 ng). **(i)**Quantitation of the single MO injections in panels b-d. **(j)**Quantitation of the combined MO injections in panels e-h. All images are lateral views, rostral to the left. Scale bar: 100 μm (b). All graphs are analyzed by t-test and the mean ± SD is shown.

